# High-throughput simulations indicate feasibility of navigation by familiarity with a local sensor such as scorpion pectines

**DOI:** 10.1101/2020.06.17.156612

**Authors:** Albert Musaelian, Douglas D. Gaffin

## Abstract

Scorpions have arguably the most elaborate “tongues” on the planet: two paired ventral combs, called pectines, that are covered in thousands of chemo-tactile peg sensilla and that sweep the ground as the animal walks. Males use their pectines to detect female pheromones during the mating season, but females have pectines too: What additional purpose must the pectines serve? Why are there so many pegs? We take a computational approach to test the hypothesis that scorpions use their pectines to navigate by chemo-textural familiarity in a manner analogous to the visual navigation-by-scene-familiarity hypothesis for hymenopteran insects. We have developed a general model of navigation by familiarity with a local sensor and have chosen a range of plausible parameters for it based on the existing behavioral, physiological, morphological, and neurological understanding of the pectines. Similarly, we constructed virtual environments based on the scorpion’s native sand habitat. Using a novel methodology of highly parallel high-throughput simulations, we comprehensively tested 2160 combinations of sensory and environmental properties in each of 24 different situations, giving a total of 51,840 trials. Our results show that navigation by familiarity with a local pectine-like sensor is feasible. Further, they suggest a subtle interplay between “complexity” and “continuity” in navigation by familiarity and give the surprising result that more complexity — more detail and more information — is not always better for navigational performance.

**Author summary:** Scorpions’ pectines are intricate taste-and-touch sensory appendages that brush the ground as the animal walks. Pectines are involved in detecting pheromones, but their exquisite complexity — a pair of pectines can have around 100,000 sensory neurons — suggests that they do more. One hypothesis is “Navigation by Scene Familiarity,” which explains how bees and ants use their compound eyes to navigate home: the insect visually scans side to side as it moves, compares what it sees to scenes learned along a training path, and moves in the direction that looks most familiar. We propose that the scorpions’ pectines can be used to navigate similarly: instead of looking around, they sweep side to side sensing local chemical and textural information. We crafted simulated scorpions based on current understanding of the pectines and tested their navigational performance in virtual versions of the animals’ sandy habitat. Using a supercomputer, we varied nine environmental, sensory, and situational properties and ran a total of 51,840 trials of simulated navigation. We showed that navigation by familiarity with a local sensor like the pectines *is* feasible. Surprisingly, we also found that having a more detailed landscape and/or a more sensitive sensor is not always optimal.

## Introduction

Scorpions present at least two unanswered scientific mysteries: the selection pressure that maintains their pectines — two ornate, sensitive, and energetically expensive sensory organs on their underbellies — and the mechanism by which they return to their burrows after emerging to hunt. We propose that these two questions are connected and that the pectines are used to navigate by a process akin to familiarity navigation, as has been previously suggested for hymenopteran insects [1].

The pectines of scorpions are paired comb-like appendages that hang mid-body from the bottom of all scorpions (Figure 1A). These organs are unique in the animal kingdom and can be traced to fossilized aquatic scorpions from the Devonian [2].

**Fig 1.**
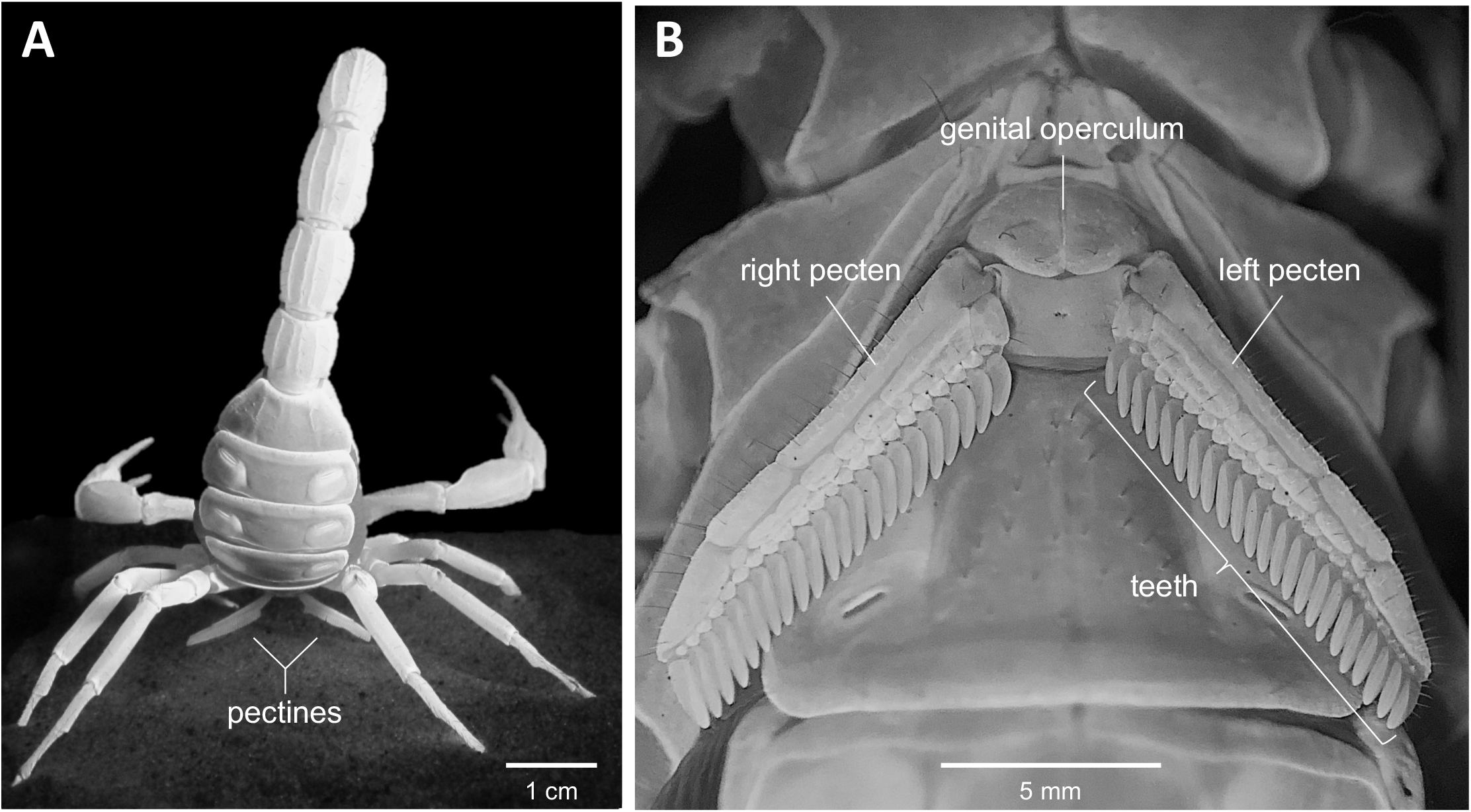
Scorpion pectines. **A**: Posterior view of a female *Hadrurus arizonensis* showing the mid-ventral pectines sweeping the substrate beneath the animal (photo by M. Hoefnagels). **B**: Ventral view of pectines of female *Paruroctonus utahensis* (photo by K. Ashford).

Behavioral [3–5], morphological [6–11], and physiological [12–15] evidence indicates that the pectines are organs of taste and touch. However, the full function of these organs remains enigmatic. In some species, the pectines are sexually dimorphic and are used by males to track putative ground-borne female pheromones [3, 16, 17]. But females have pectines too, and in many species the organs are not strongly dimorphic. Together, these observations raise the question: What else could the pectines be doing? Here we expand on our recent hypothesis that the pectines may guide navigation by a process of chemo-textural familiarity [18].

### Scorpion pectines

Understanding this hypothesis requires some background in pecten morphology and physiology. The flexible spine of each pecten extends ventro-laterally from mesosomal segment two (Figure 1A) and supports a species-dependent number of stout, ground-facing teeth (Figure 1B). The distal face of each tooth contains a dense array of minute, peg-shaped sensilla called “pegs” (Figure 2). In some species, such as male *Smeringerus mesaensis* (formerly *Paruroctonus mesaensis*), the number of pegs per tooth reaches 1600 for a total of over 100,000 across both pectines [4, 19]. The peg slits are oriented perpendicular to the travel of the ground as the teeth fan out when the pectines brush the surface.

**Fig 2.**
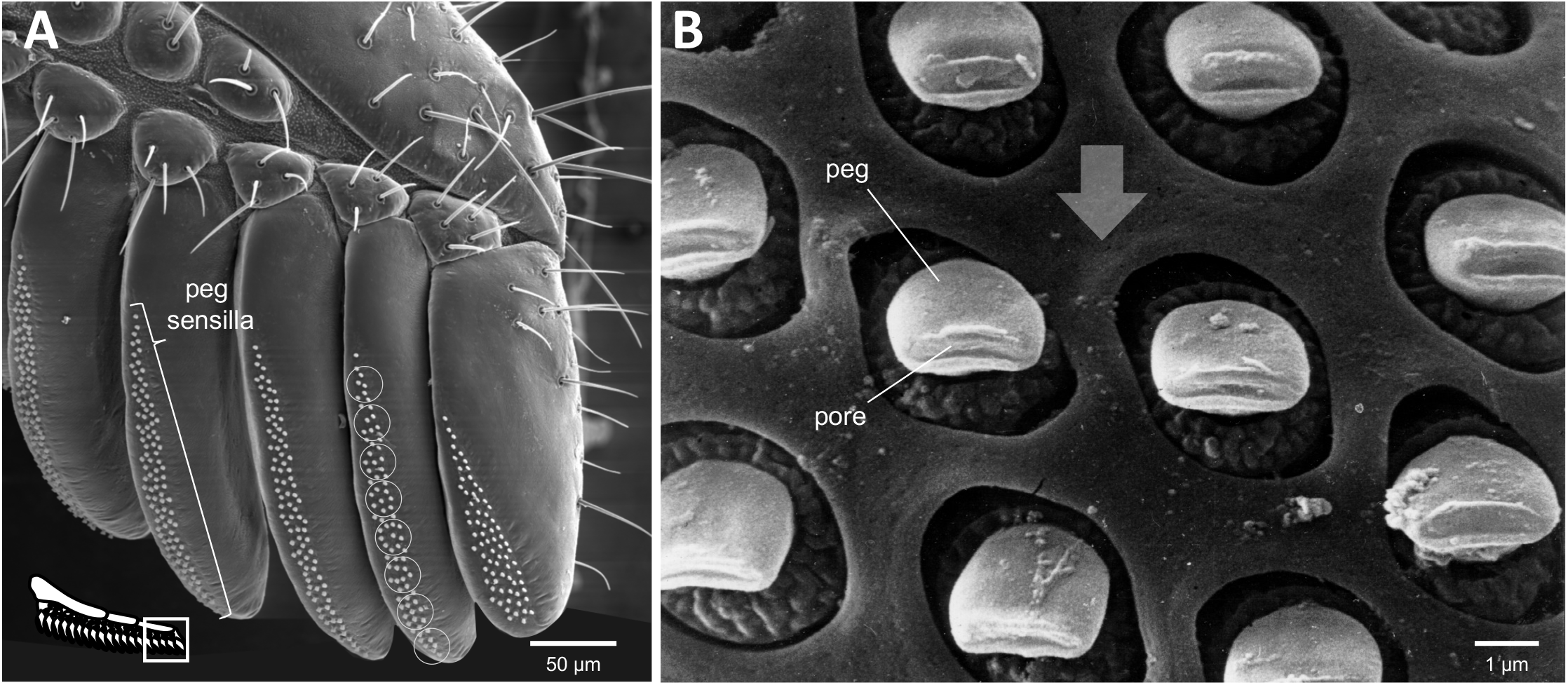
Chemo-tactile peg sensilla. **A**: An SEM of the five distal teeth of a male *P. utahensis* pecten shows the patches of peg sensilla that coat the ground-facing surfaces of each tooth (photo by E. Knowlton). The superimposed circles on the penultimate tooth show the approximate resolution of chemical discrimination based on peg response patterns and pecten sweep kinetics (see text). **B**: An expanded view of a patch of peg sensilla from male *Smeringerus mesaensis* shows the consistent orientation of their pore tips perpendicular to the relative movement of the ground (arrow indicates direction of ground movement).

Each peg is supplied with about a dozen sensory neurons. All but one of these neurons have unbranched dendritic outer segments that extend into a fluid-filled chamber just proximal to the peg’s slit-shaped terminal pore [7, 8] and can best be classified as gustatory neurons [20]. The remaining neuron terminates in a tubular body near the peg base, which is a hallmark of arthropod mechanoreceptors [8, 21]. The peg neurons course along the pecten spine and through the pectinal nerve to the subesophageal ganglion (SEG) in the posterior brain [10, 11, 19, 22, 23]. A cross-section of the pectinal neuropil in the SEG shows an ordered, topographical arrangement of sensory input based on the position of the teeth on the pectinal spine [24].

Extracellular, electrophysiological recordings show that peg neurons respond to a variety of near-range, volatile organics including alkanes, alcohols, aldehydes, ketones, and esters, especially in the 6-10 carbon chain length [12]. Deflection of the peg while recording from the base also elicits a phasic mechanoreceptor response [12, 25]. Tip recordings from hundreds of *P. utahensis* pegs with citric acid, ethanol, or KCl added to the electrolyte in the glass electrode show that the pegs respond to all three stimulants in a characteristic yet uniform way [15, 26].

The arrangement of the peg arrays also appears to increase information resolution and acquisition speed. In isolation, and within the time constraints of normal pectinal behavior, individual pegs are unable to discern a chemical’s identity [15]. However, integrating information simultaneously from multiple pegs yields precise information that is in line with the short duration of a typical pectinal sweep [14]. In particular, it was found that a minimum of eight pegs working in parallel were required to distinguish citric acid from ethanol [15]. The superimposed circles on the pectinal tooth in Figure 2A gives a sense of this putative spatial resolution.

### Navigation by familiarity

The *Navigation by Scene Familiarity Hypothesis* is a provocative and elegant idea that has been proposed in studies of the visual navigation of hymenopteran insects. It suggests that the dense arrays of ommatidia in insect compound eyes are used to transform scenes from the insect’s visual world into spatial matrices of information. To navigate back to a hive or food source, the insect simply moves in the direction that yields visual scenes that are the most similar to the scenes it learned during a previous training excursion [1, 27, 28]. In other words, the insect navigates back by choosing a direction that maximizes “familiarity.” The hypothesis appears congruent with numerous behavioral observations [29–36]. Furthermore, there are now robots and simulated agents that navigate autonomously via familiarity [1, 18, 37–39], and some studies [40, 41] have suggested how neural tissue, such as the central complex and mushroom bodies [42–51], might be organized to accommodate familiarity-based navigation.

### Navigation by familiarity with a local sensor

In this paper, we consider the hypothesis that the dense fields of peg sensilla on pectines are analogous to the tightly packed ommatidia in compound eyes, detecting matrices of chemical and textural information that are used for navigation by familiarity. While our hypothesis is inspired by visual familiarity navigation, the pectines are a sensor that only senses the local environment, which is critically different from a non-local sensor like eyes that gather information from both near and far.

This work uses computer simulations to establish whether and when it is possible to navigate by familiarity using a local sensor. Throughout this manuscript, we will use NFLS as our acronym for *Navigation by Familiarity with a Local Sensor*. We have developed a general computer model of NFLS and use a broad base of morphological, physiological, and behavioral information to configure a simulated sensor that models the pectines. Previous work has suggested a subtle interplay between the environmental and sensory conditions needed to enable familiarity navigation [18, 37, 52]. To explore this interplay and the impact of different parameters on the performance of NFLS, we comprehensively simulated 51, 840 different unique combinations of sensory, environmental, and situational properties.

## Models

### Computational model of NFLS

Our computational model (Figure 3) depends on a number of simplifying assumptions:

1. The agent’s environment does not change during or between learning walks and navigation. This is plausible because scorpions are nocturnal and the desert winds are calm at night, especially on the leeward side of the dunes (pers. obs.; [53]).
2. The shape of the agent’s sensor is a constant rectangle — in scorpion terms, the two pectines move together. Although experimental data indicate that the pectines move independently (see Figure 3 in S1 Appendix), the relevant consideration for NFLS is only whether both pectines achieve the same orientation within a short amount of time. This requirement could be satisfied by a wide variety of pecten movement patterns, but we chose to model the simplest.
3. There is a meaningful minimum and maximum stimulus. Just as humans experience nothing but pure darkness or blinding white outside of a certain range of light intensity, we assume that the agent’s experience of chemical concentration or mechanical pressure clips to some range.

**Fig 3.**
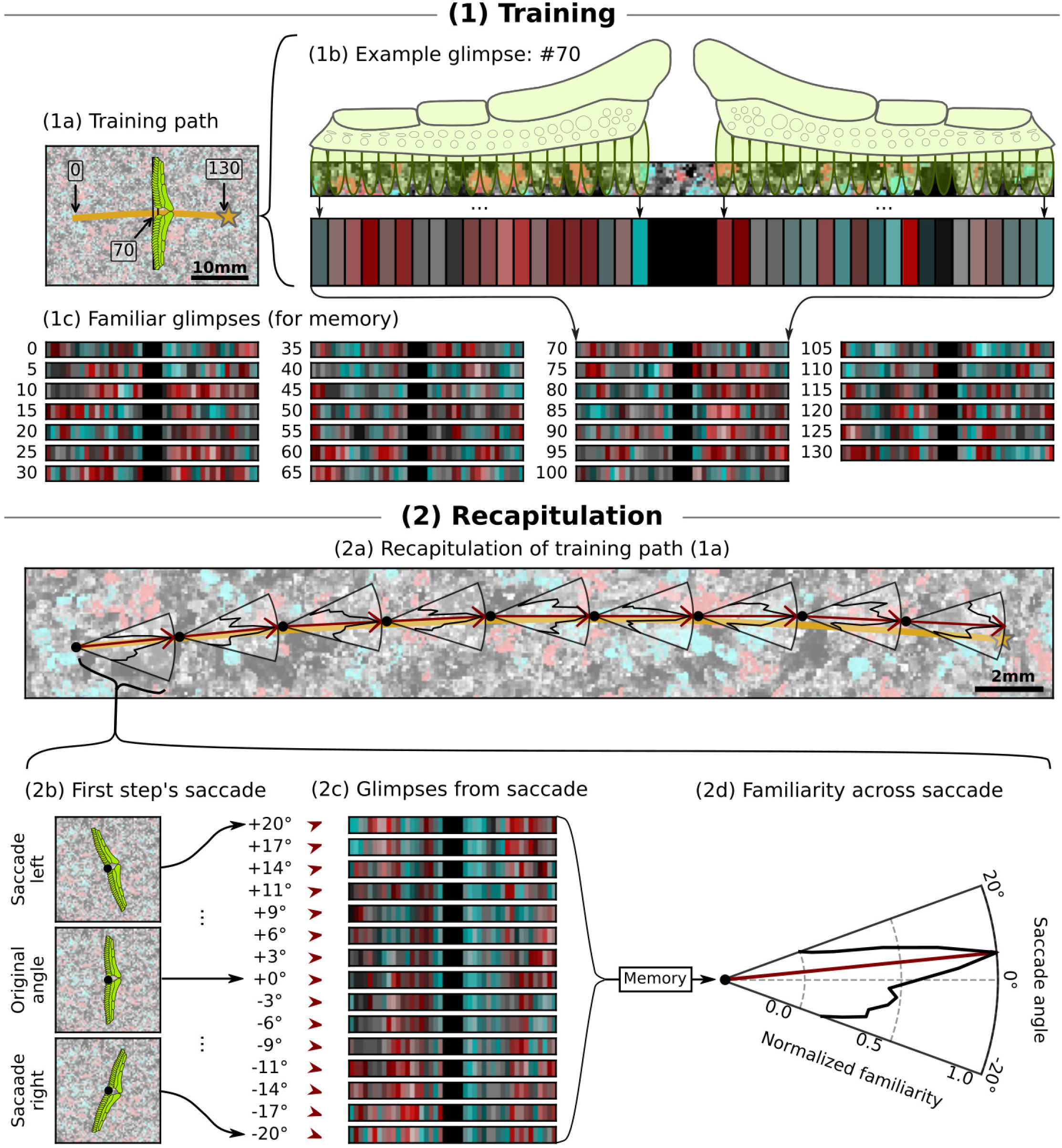
Schematic illustration of NFLS. **(1a)** shows the training path in gold. **(1b)** shows how the model constructs a specific glimpse from the landscape. **(1c)** shows some of the familiar glimpses gathered along the training path. **(2a)** Each red arrow indicates a step; each wedge is a normalized polar plot of familiarity as a function of saccade angle. **(2b)** shows the saccading of the pectines during the first step; **(2c)** shows the resulting glimpses, and **(2d)** shows the resulting familiarity “compass.” See section 4 in S1 Appendix for the parameters of the shown agent.

### Sensing the environment

The landscape in which the agent navigates is represented by a two-dimensional 8-bit hue-saturation-brightness (HSB) color image. The image is interpreted as a view from above. The brightness channel represents height and thereby texture; higher values represent greater height. The hue channel represents the dominant chemistry of each pixel; each of the 255 possible values represents a distinct chemical. The saturation channel represents, at each pixel, the concentration of the chemical indicated by its hue.

The agent’s sensor is represented as a grid of large sensor “pixels” of equal rectangular shape. The dimensions of the sensor pixels are given in terms of landscape pixels. A pecten-like sensor could, for example, be a 40 × 1 grid of sensor pixels that each represents a tooth and corresponds to a 6 × 12 area on the landscape. The choice to model the sensor as a grid corresponds to the grid-like physiology and neurology of the pectines [10, 11, 15].

We call the grid of sensor pixels resulting from the agent’s using its sensor to observe the landscape a “glimpse” consistently with the familiarity literature. Despite the vision-specific connotations of the word in common use, in this paper a “glimpse” consists of chemo-textural and not visual information. When referring to visual information, we use the word “scene.” We denote the sensor matrix glimpsed by the agent at position (*x, y*) and orientation *θ* by Glimpse(*x, y, θ*).

To compute Glimpse(*x, y, θ*), we first crop the landscape to the rectangle representing the sensor centered on (*x, y*) and rotate to the agent’s orientation given by *θ*. (Rotations use nearest-neighbor interpolation.) The resulting image represents the relevant portion of the landscape that is directly under the sensor. The grid of sensor pixels is then superimposed on the cropped portion of the landscape. The value of each sensor pixel is computed from the block of landscape pixels it covers: The brightness of the sensor pixel is set to the mean brightness of the block, the chemical identity of the sensor pixel is set to the chemical that has the greatest total concentration across the block, and the concentration is set to the total concentration of the chosen chemical. Note that each sensor pixel detects only the chemicals directly in contact with it. This restriction is justified by [14], which shows that the response of peg sensilla to pure volatiles, such as 6-carbon alcohols, falls to essentially nothing when the stimulant is further than 10 microns away.

The resulting sensor matrix is itself an 8-bit HSB image. To simulate a less sensitive sensor, the brightness values in the sensor matrix are each rounded to the closest of a small number of valid levels. If the number of allowed levels is set to two, for example, the result is a binary sensor: each pixel’s brightness is rounded to the nearer of 0 or 255. Our model can also independently apply similar rounding to the hue and saturation in order to simulate a configurable sensitivity to chemical differences, but we have not done so in this work.

### Training

Given a training path as a sequence of points (*x*_1_*, y*_1_), (*x*_2_*, y*_2_)*, ...,* (*x*_*n*_, *y*_*n*_) we collect the training glimpses *T*_1_*, T*_2_*, ..., T*_*n*−1_ as

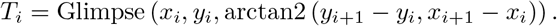

Training is shown schematically in panel 1 of Figure 3.

### Navigation

Following our own previous work and all prior work we are aware of in the literature [1, 28, 40], we discretize the process of navigation by familiarity into small spatial movements we call “steps.” Steps in our model do not correspond to any particular movement of the scorpion’s legs: rather, they are a necessary construct for practically modeling familiarity navigation. In our model, at each step the agent rotates its sensor across a range of orientations; at each orientation, it tests the familiarity of the resulting glimpse of the environment. The agent then chooses the direction that was most familiar and moves by a small fixed step length in that direction. This back-and-forth rotation, called “saccading,” is observed in other animals, such as ants and bees, hypothesized to use familiarity navigation [54, 55]. Saccading allows the animal to test a range of candidate directions in which it could move. Rather than rotating their entire body, like bees do, we propose that scorpions saccade by moving their pectines, which are highly mobile when the animal walks (see, for example, Figure 3 in S1 Appendix). The process of saccading and choosing an orientation is shown schematically in panel 2 of Figure 3.

To complete a step, the set of saccade offsets *S* is first defined as the *S*_n_ angles evenly distributed over the interval 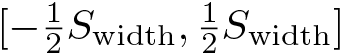 including the endpoints, where *S*_width_ is the saccade width parameter and *S*_*n*_ is a parameter denoting the number of glimpses per step. If the agent is currently at position (*x, y, θ*), its new heading is chosen as:

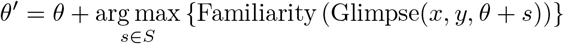

The agent then steps in the chosen direction, achieving a new position of

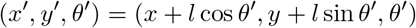

where *l* is the step length parameter.

We define the familiarity of a sensory glimpse *G* as

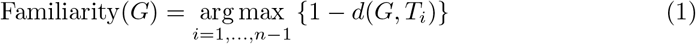

where *d*(·,·) ∈ [0, 1] is a metric (in the informal rather than strict sense) among glimpses. We define *d*(·, ·) as a weighted combination of a metric for textural differences and a metric for chemical differences:

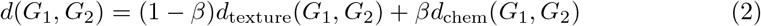

where *β* ∈ [0, 1] is the weight that determines how much relative importance to give to chemistry versus texture. For the texture difference metric, we use the <u>s</u>um of <u>a</u>bsolute <u>d</u>ifference<u>s</u> (SADS) metric [52]:

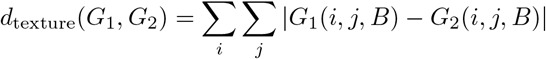

where *G*(*i, j,* {*H, S,* or *B*}) denotes the H, S, or B component of the (*i, j*)th sensor pixel in glimpse *G*. For the chemical difference metric, we define:

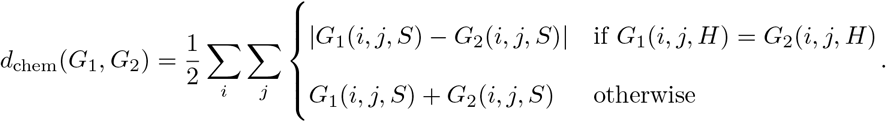

In words, when the (*i, j*)th sensor pixels in *G*_1_ and *G*_2_ have the same chemical identity, the summand is the difference between their concentrations. When they have different chemical identities, the summand is the sum of their concentrations. This yields the intuitive result that there is a bigger difference between a high concentration of chemical A and a high concentration of chemical B than there is between a high concentration of A and a low concentration of B.

This method of comparing a glimpse to all memorized training glimpses (Equation 1) corresponds to the perfect recall of training glimpses previously explored for non-local sensors in [1, 37, 52]. In an effort to increase realism, some researchers [1, 28, 40] have trained general and biologically-inspired artificial neural networks to approximate the familiarity function. For our work, however, neural networks would introduce a noisy black box that could distort or overwhelm the effects of other variables and mask results.

The code for our model and visualizations is open-source and publicly available [56]; details on its optimized implementation can be found in section 4 of S1 Appendix.

### High-throughput simulated navigation trials

Using cluster computing we conducted a total of 51,840 trials of simulated NFLS, each with a unique combination of sensory, environmental, and situational parameters. The trials were constructed as every possible combination of the tested values for each parameter, which are listed below. This high-throughput highly parallel methodology is primarily inspired by high-throughput virtual screening and optimization efforts in materials science such as [57] and [58]. Similar methods have also attained prominence in proteomics, computational chemistry, pharmacology, and many other fields.

Trials were independently run in a parallel fashion, with each trial ending when its agent (1) reached the end of the training path, (2) wandered more than 450 px = 33 mm away from the training path, or (3) exceeded the maximum number of navigation steps, which we set to three times the number of steps that would be necessary to recapitulate the path without detours. We considered a trial that reached the end of the path to be a successful navigation. (A real-world agent could detect that it had reached its target through tactile, chemical, or even familiarity-based cues as proposed in [1].) Trials where the agent wanders too far from the training path were terminated to prevent the agent from exiting the landscape. By fixing a constant maximum distance rather than terminating when the agent exited the landscape, we avoided biasing our results by giving agents that got lost further from the edge of the landscape a greater chance of recovering. Since the likelihood that the agent would re-encounter the path strongly decreased with distance from the path, it is very unlikely that this condition terminated any trial that would have otherwise succeeded. Finally, by fixing a maximum number of steps, we placed a firm upper bound on the duration of each trial. Of the few trials that were terminated for this reason, most had extreme parameters, and the agent had become stuck in a loop which indicates that this constraint did not meaningfully bias our results (see section 6 in S1 Appendix).

### Tested Parameters

We chose our parameters to represent a broad range of plausible hypotheses about scorpions, their pectines, and their environments. Whenever possible, behavioral, physiological, neurological, and chemical data were used to guide parameter choice.

*Environmental properties —* **Landscape** We stitched together landscapes from grey-scale microscope images of desert sand (Figure 4). The brightness of each pixel roughly corresponds to its height, encoding the texture of the sand. Chemical information, simulated by hue, is added randomly to contiguous areas of greatest height in each trial’s landscape in a manner controlled by the next two parameters. Details of the generation of the landscapes and the addition of the chemical information can be found in sections 1 and 2 of S1 Appendix.

**Fig 4.**
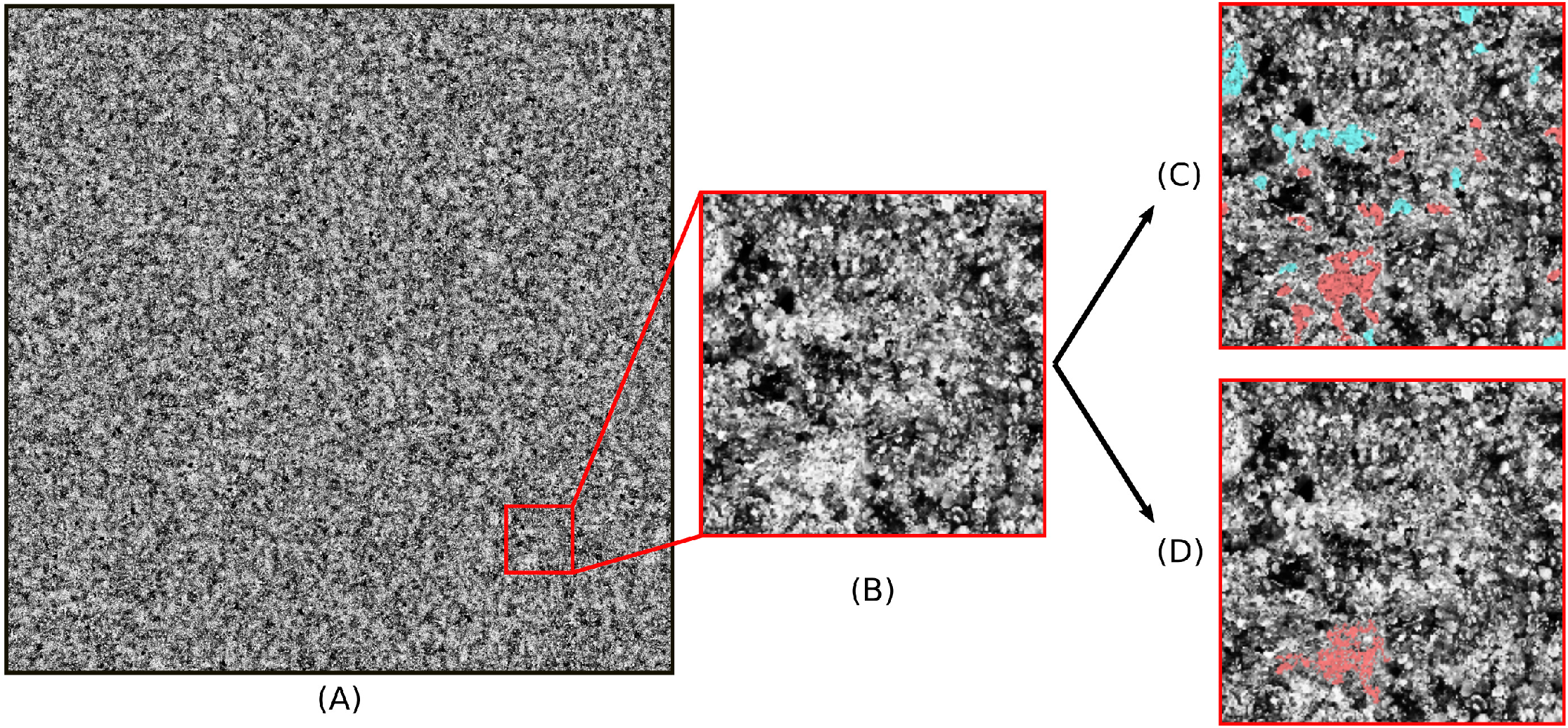
Example landscape. **(A):** The first of the four landscapes generated. **(B):** Detail of (A). **(C):** Same detail of (A) with chemistry added, using two chemicals and a chemical scale of 0.5 mm. **(D):** Same detail of (A) with chemistry added using a chemical scale of 2.0 mm.

Each landscape is 2000 × 2000px at a resolution of 13.6 px/mm, giving a side length of 14.7cm.

<u>Tested Values:</u> Four unique landscapes randomly generated in this manner.

**Number of distinct chemicals** <u>Tested Values:</u> 1, 2, 4.

**Spatial scale of chemical detail** The minimum size that a contiguous area of greatest height in the landscape must have to be colored, which represents saturating it with some chemical. The chemical information added to the landscape becomes coarser and sparser as this parameter increases. Technical details are given in section 2 of S1 Appendix.

<u>Tested Values:</u> 0.5mm, 1.0mm, 2.0mm.

*Sensor and agent properties —* **Step size** We fix a step size of 1.3mm that aligns well with the time resolution of the experimental data from [3] that we used to estimate other agent parameters (see section 3 in S1 Appendix).

<u>Tested Value:</u> 1.3mm = 17.72 px.

**Spatial resolution** We base our simulated sensor on the pectines of female *P. utahensis*, which have about 40 teeth across the two pectines [59]. Each sensory resolution represents a different hypothesis about the granularity with which the scorpion’s brain distinguishes between the pegs on a tooth. For example, 40 × 1 represents a situation where all the pegs on each tooth offer a single aggregate stimulus to the brain, while 40 × 6 represents a scenario where independent stimuli from six different groups of pegs on each tooth are processed at the brain.

The grouped nature of the pegs is supported by experimental evidence: It takes approximately eight pegs working in tandem to distinguish between chemicals [15]. While each peg contains a mechanoreceptor [8, 9], and the kinetics of the mechanosensory response [12] may yield finer-grained resolution, the textural discrimination power of the pegs has not been tested. As such, we chose to use the more conservative values based on the chemosensory response.

Regardless of its resolution, the sensor covers a 19.5mm × 0.9mm area, equal to the typical bounding box of the teeth of the pectines of female *P. utahensis*. The middle 2mm × 0.9mm of the sensor, corresponding to the approximately 2mm of space between the pectines (Figure 1), is blacked out.

<u>Tested Values:</u> 40 × 1, 40 × 2, 40 × 4, 40 × 6.

**Number of texture levels** Each value represents a hypothesis as to the number of distinct degrees of flexion that peg mechanoreceptors can distinguish. At two levels of intensity, texture values will be rounded to the nearer of 0 or 255; at 4, the nearest of 0, 64, 127, and 255, and so on.

<u>Tested Values:</u> 2, 4, 8, 16.

**Weighting of familiarity between texture and chemistry** The value of *β* in Equation 2. At *β* = 0.0, only texture is considered for familiarity. At *β* = 1.0, only chemical information is considered.

<u>Tested Values:</u> 0.0, 0.25, 0.5, 0.75, 1.0.

**Saccade width** We used behavioral data from [60] to estimate the range of orientations achieved by the pectines within a single 1.3mm step (details can be found in section 3 of S1 Appendix).

<u>Tested Values:</u> 30°, 40°, 50°.

**Number of glimpses per step** The scorpion’s pectines sweep over the ground, sending a continuous stream of neural signals to the brain. Existing experimental work on the neurobiology of the pectines, however, does not yield a clear picture of their response time or temporal dynamics. As a result we chose to guide our choice using the general arachnid literature. In [61] the minimum response time of a spider mechanoreceptor is estimated at 1.2 ms. Our analysis of behavioral data (section 3 in S1 Appendix) led us to estimate that the scorpion takes about 180ms to move 1.3mm, the length of one step. Dividing yields a maximum of approximately 156 glimpses per step.

To be conservative and working under the assumption that the chemical response time of the pegs is likely slower, we cut this upper bound by about an order of magnitude and chose 25 glimpses/step, yielding an angular resolution between 1.2° and 2°, agreeing with typical literature values of 1° to 5° [1, 28, 40].

<u>Tested Value:</u> 25 glimpses/step.

*Situational properties —* **Start offset** We test navigation both from the beginning of the training path and from a point offset from it. The offset is given as a counterclockwise change to the agent’s orientation and a translation given as a percentage of the sensor’s width. For example, if the agent’s original position is (*x, y, θ*) and the offset is 10 % at −7°, then the offset position is

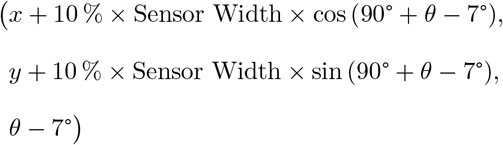

<u>Tested Values:</u> None, 10 % at −7°, −10 % at 7°.

**Training path** We define diagonal S-shaped training paths with various curvatures *α* ∈ [0, 1] given by the curve:

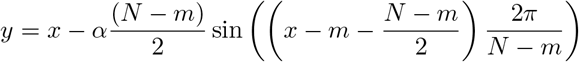

for *x* ∈ [*m, N* − *m*] where *N* = 2000 px is the landscape size and *m* = 500 is the margin from the edge of the landscape. Each path is then cropped equally at the beginning and end until its length is equal to 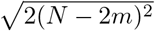, which is the length of the shortest possible path (*α* = 0). This prevents training path lengths from varying and unintentionally influencing navigational performance.

We chose to consider only training paths with low curvature. A path of high curvature can be approximated by a series of linear steps only if those steps are sufficiently small. Training paths of high curvature would thus have artificially low navigational success due to this geometric effect. These failures would not be scientifically relevant, since, as mentioned earlier in the description of the model, the discretization of the navigation process is a concession to practical modeling: Actual scorpions move their pectines and change direction with sufficient frequency to follow a path that is highly curved at any physically relevant scale.

Our choice to focus on low curvature paths also agrees with a prominent hypothesis — path integration [1, 62–64] — for how animals that navigate by familiarity could conduct their training walks. Path integration results in straight line training paths. It has also been demonstrated in spiders, another arachnid [65–67].

<u>Tested Values:</u> *α* = 0.1, 0.2.

### Success metrics

We defined and computed five metrics for every trial:
 
1. **Success**: Whether the agent reached the end of the training path;
2. **Path completion**: The furthest point on the path where the agent traversed at least 5 % of the training path’s length consecutively;
3. **Path coverage**: The percentage of the training path the agent traversed;
4. **Number of captures**: How many times the agent encountered the training path and subsequently traversed at least 5 % of its length;
5. **Root-mean-square deviation**: The root-mean-square deviation (RMSD) of the agent’s position from the training path over the entire trial.

## Results and Discussion

Hereafter, we define a “sensory configuration” as a unique combination of values for the parameters listed as sensory properties above. An “environmental configuration” is defined similarly from the list of environmental properties. A “sensory-environmental configuration” is a unique combination of one of each. We define a “situation” as a unique combination of a start offset, a training path curvature, and a specific landscape (as mentioned above we tested four different but identically generated landscapes). Every possible sensory-environmental configuration was tested in all situations. In other words, trials for every possible pairing of a sensory and environmental configuration were run.

### Most successful configurations

For each sensory, environmental, and sensory-environmental configuration, we computed its success rate as the percentage of successful trials among all that used the given configuration. The five most successful sensory, environmental, and sensory-environmental configurations are given in Table 1, Table 2, and Table 3 respectively. We computed the success rates twice: first across trials without start offsets and then across all trials. The first calculation serves to establish the fundamental limitations of NFLS in an ideal world, while the second introduces realistic imperfections.

**Table 1.** Top sensory configurations by success rate.

**Over trials without start offsets (72 trials per configuration)**
| % successful | # texture levels | Sensory configuration |  |  | Avg. properties of successes |  |  |
| --- | --- | --- | --- | --- | --- | --- | --- |
|  |  | Saccade | Resolution | Chem. weight | Coverage | Captures | RMSD |
| 100 % | 16 | 30° | 40 × 4 | 0.0 | 95 ± 5 % | 1.4 ± 0.5 | 0.37 ± 0.38 mm |
| 90 % | 4 | 40° | 40 × 4 | 0.2 | 98 ± 0 % | 1.0 ± 0.2 | 0.16 ± 0.04 mm |
| 88 % | 8 | 30° | 40 × 4 | 0.0 | 96 ± 6 % | 1.3 ± 0.7 | 0.26 ± 0.33 mm |
| 88 % | 16 | 30° | 40 × 6 | 0.0 | 96 ± 4 % | 1.3 ± 0.5 | 0.31 ± 0.36 mm |
| 88 % | 16 | 50° | 40 × 4 | 0.0 | 92 ± 15 % | 1.0 ± 0.0 | 1.26 ± 2.71 mm |

**Over all trials (216 trials per configuration)**
| % successful | # texture levels | Sensory configuration |  |  | Avg. properties of successes |  |  |
| --- | --- | --- | --- | --- | --- | --- | --- |
|  |  | Saccade | Resolution | Chem. weight | Coverage | Captures | RMSD |
| 58 % | 4 | 50° | 40 × 4 | 0.0 | 93 ± 12 % | 1.0 ± 0.0 | 0.37 ± 0.44 mm |
| 58 % | 16 | 40° | 40 × 6 | 0.0 | 94 ± 10 % | 1.1 ± 0.3 | 0.66 ± 1.52 mm |
| 54 % | 8 | 40° | 40 × 6 | 0.0 | 95 ± 5 % | 1.1 ± 0.3 | 0.29 ± 0.15 mm |
| 54 % | 16 | 30° | 40 × 4 | 0.0 | 95 ± 4 % | 1.2 ± 0.4 | 0.35 ± 0.30 mm |
| 52 % | 16 | 40° | 40 × 6 | 0.2 | 91 ± 20 % | 1.0 ± 0.3 | 0.76 ± 1.97 mm |

“% successful” indicates the percentage of all trials with the given sensory configuration that were successful. “Avg. properties of successes” show averages over all trials with the given sensory configuration that were successful; the values given after ± are standard deviations.

**Table 2.** Top environmental configurations by success rate.

**Over trials without start offsets (1920 trials per configuration)**
| % success | Environmental configuration |  | Avg. properties of successes |  |  |
| --- | --- | --- | --- | --- | --- |
|  | # chems. | Chem. scale | Coverage | Captures | RMSD |
| 48 % | 1 | 2.00 mm | $97 \pm 8 \%$ | $1.1 \pm 0.3$ | $0.38 \pm 1.31$ mm |
| 47 % | 4 | 2.00 mm | $97 \pm 8 \%$ | $1.1 \pm 0.3$ | $0.39 \pm 1.36$ mm |
| 47 % | 2 | 2.00 mm | $96 \pm 8 \%$ | $1.1 \pm 0.3$ | $0.40 \pm 1.42$ mm |
| 47 % | 4 | 0.50 mm | $96 \pm 10 \%$ | $1.1 \pm 0.3$ | $0.43 \pm 1.46$ mm |
| 46 % | 2 | 0.50 mm | $96 \pm 10 \%$ | $1.1 \pm 0.3$ | $0.51 \pm 1.75$ mm |

**Over all trials (5760 trials per configuration)**
| % success | Environmental configuration |  | Avg. properties of successes |  |  |
| --- | --- | --- | --- | --- | --- |
|  | # chems. | Chem. scale | Coverage | Captures | RMSD |
| 27 % | 1 | 2.00 mm | $92 \pm 15 \%$ | $1.1 \pm 0.3$ | $0.85 \pm 2.28$ mm |
| 26 % | 4 | 2.00 mm | $92 \pm 15 \%$ | $1.1 \pm 0.3$ | $0.82 \pm 2.28$ mm |
| 26 % | 2 | 2.00 mm | $92 \pm 16 \%$ | $1.1 \pm 0.3$ | $0.83 \pm 2.31$ mm |
| 25 % | 4 | 0.50 mm | $90 \pm 19 \%$ | $1.1 \pm 0.4$ | $1.04 \pm 2.67$ mm |
| 24 % | 2 | 0.50 mm | $90 \pm 19 \%$ | $1.1 \pm 0.3$ | $1.10 \pm 2.78$ mm |

“% successful” indicates the percentage of all trials with the given environmental configuration that were successful. “Avg. properties of successes” show averages over all trials with the given environmental configuration that were successful; the values given after ± are standard deviations.

**Table 3.** Top sensory-environmental configurations by success rate.

**Over trials without start offsets (8 trials per configuration)**
| % succ. | Env. configuration |  | Sensory configuration |  |  |  | Avg. properties of successes |  |  |
| --- | --- | --- | --- | --- | --- | --- | --- | --- | --- |
|  | Chems. | Chem. scale | Texture lvls. | Saccade | Res. | Chem. | Coverage | Captures | RMSD |
| 100 % | 1 | 0.50 mm | 16 | 30° | $40 \times 4$ | 0.0 | $95 \pm 5 \%$ | $1.4 \pm 0.5$ | $0.37 \pm 0.38$ mm |
| 100 % | 1 | 0.50 mm | 2 | 30° | $40 \times 4$ | 0.5 | $98 \pm 0 \%$ | $1.0 \pm 0.0$ | $0.15 \pm 0.04$ mm |
| 100 % | 1 | 0.50 mm | 4 | 30° | $40 \times 2$ | 0.5 | $98 \pm 0 \%$ | $1.0 \pm 0.0$ | $0.15 \pm 0.03$ mm |
| 100 % | 2 | 0.50 mm | 2 | 30° | $40 \times 6$ | 0.5 | $98 \pm 0 \%$ | $1.0 \pm 0.0$ | $0.15 \pm 0.04$ mm |
| 100 % | 2 | 0.50 mm | 2 | 30° | $40 \times 6$ | 0.8 | $98 \pm 0 \%$ | $1.0 \pm 0.0$ | $0.16 \pm 0.03$ mm |

**Over all trials (24 trials per configuration)**
| % succ. | Env. configuration |  | Sensory configuration |  |  |  | Avg. properties of successes |  |  |
| --- | --- | --- | --- | --- | --- | --- | --- | --- | --- |
|  | Chems. | Chem. scale | Texture lvls. | Saccade | Res. | Chem. | Coverage | Captures | RMSD |
| 67 % | 1 | 1.00 mm | 16 | 40° | $40 \times 6$ | 0.2 | $93 \pm 7 \%$ | $1.1 \pm 0.2$ | $0.38 \pm 0.32$ mm |
| 62 % | 1 | 0.50 mm | 2 | 30° | $40 \times 4$ | 0.5 | $88 \pm 24 \%$ | $0.9 \pm 0.2$ | $1.41 \pm 3.91$ mm |
| 62 % | 1 | 0.50 mm | 2 | 40° | $40 \times 6$ | 0.5 | $85 \pm 27 \%$ | $1.1 \pm 0.5$ | $1.55 \pm 3.33$ mm |
| 62 % | 2 | 1.00 mm | 16 | 40° | $40 \times 6$ | 0.2 | $95 \pm 4 \%$ | $1.1 \pm 0.2$ | $0.31 \pm 0.24$ mm |
| 62 % | 1 | 2.00 mm | 16 | 40° | $40 \times 6$ | 0.5 | $89 \pm 24 \%$ | $0.9 \pm 0.2$ | $1.72 \pm 4.30$ mm |

“% successful” indicates the percentage of all trials with the given sensory-environmental configuration that were successful. “Avg. properties of successes” show averages over all trials with the given sensory-environmental configuration that were successful; the values given after ± are standard deviations.

### Feasibility of NFLS without offsets

Without start offsets, there is at least one physically plausible sensory configuration that succeeds in 100 % of tested environments and situations. We believe that this is strong evidence that the scorpion’s environment and physiology contain and sense sufficient information to enable NFLS. The existence of other highly successful sensory configurations — about 5 % of sensory configurations had a success rate above 80 % without offsets (Figure 6 in S1 Appendix) — further supports this conclusion. Among complete sensory-environmental configurations, 2 % (33 configurations) achieved a 100 % success rate across all situational parameters (Figure 5 in S1 Appendix).

### Feasibility of NFLS with offsets

In general, trials with offsets were three times less successful than those without. Thus, in considering all trials, we expected and saw a decrease in success rates: The success rate of the best sensory configuration declined from 100 % to 58 %, while the success rate of the top sensory-environmental configuration declined from 100 % to 67 % (Table 1, Table 3). Nonetheless, the top configurations still succeeded in many situations with offsets, which supports the feasibility of NFLS under realistic conditions.

**Fig 5.**
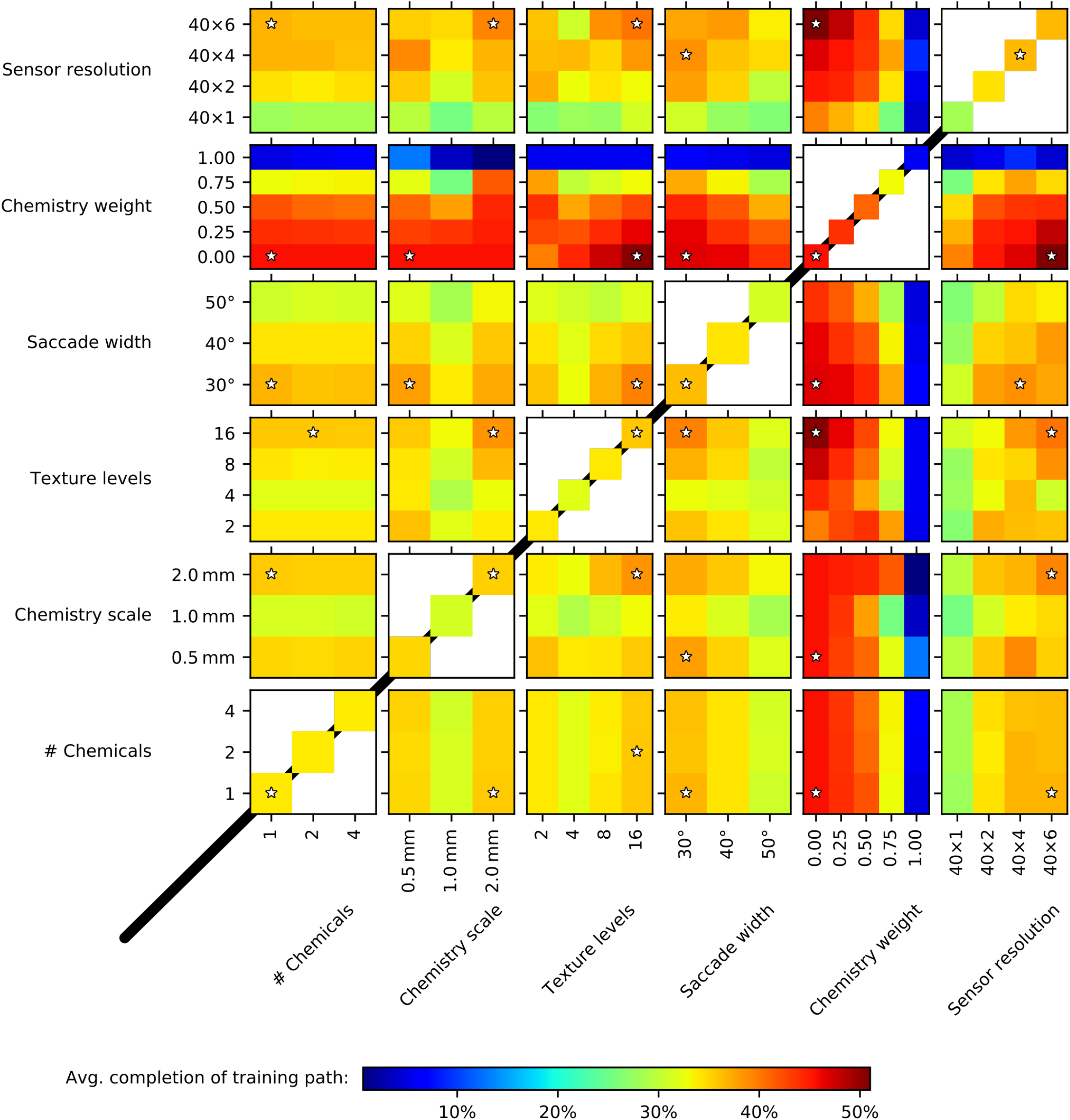
Effect of variables on navigational performance. Each heatmap shows the average training path completion for the various combinations of values across two variables. Heatmaps below the 45° black line of symmetry are mirror duplicates of those above it and are included for convenience only. The heatmaps on the diagonal that show the effect of a single variable are reproduced as line plots with error bars in figure 7 of S1 Appendix.

Further, the data clearly indicate the agent’s ability to “catch” the training path and follow it to the goal regardless of whether an offset is present. The inclusion of offset runs in our calculations does not meaningfully affect the average number of captures among successful trials. The consistently high path coverage of successful trials supports the notion that all consistently successful agents follow the path closely after encountering it regardless of whether they have an offset. Finally, repeated capture was not a typical feature of any highly successful configuration, even though in general trials with more than one capture succeeded just as frequently as trials with one: 47 % of trials with one capture succeeded while 52 %, 42 %, and 50 % of trials with 2, 3, and 4 captures, respectively, did likewise. This indicates that repeated recapture, though capable of supporting navigational success, does not do so consistently and is not a reliable mechanism for robust navigation.

As such, we believe that simple behaviors such as zig-zagging or walking out in a spiral when overall familiarity is low could dramatically increase the agent’s chances of encountering the path after starting at an offset and thus significantly increase navigational success. These behaviors have been observed in other animals hypothesized to use familiarity navigation [68, 69]; there is at present, however, no evidence for or against such behavior in scorpions. Studying the movement patterns of scorpions in their native habitats would be an important contribution to the literature and could further illuminate these issues.

### Existence of “fragile” successes

Approximately 40 % of sensory-environmental configurations were “fragile,” meaning that even across a group of favorable situations that lacked offsets and were substantively identical, the configurations succeeded neither rarely *nor* consistently: their success rates were between 20 and 60 % (see Figure 6 in S1 Appendix). Because the only differences between situations without offsets were the specific landscape and small shifts in the training paths, the successes and failures of these configurations were for all practical purposes coincidental.

A configuration for a navigational scheme is not plausible if agents using it cannot succeed across the many different situations real animals would encounter in their lives. If a configuration is fragile, it is not plausible. In order to determine, however, whether a configuration is fragile or robust, one must test it in many situations, including very similar ones. The prevalence of fragile configurations in our data illustrates the importance of using high-throughput simulations and not assuming that a few successes indicate a robust scheme. These cautions apply both to this paper and the literature more broadly: visual familiarity navigation could also have fragile successes. In the case of a visually guided agent navigating toward one of two similar trees, for example, a very small change to the environment could be sufficient to make a scene containing the wrong tree slightly more familiar than a scene containing the right one, causing the agent to get lost. Thus we conclude that many trials are needed to expose fragile successes in any work on simulated navigation and that additional simulations could usefully augment even large-scale work like ours.

### Trends in navigational performance

In addition to finding configurations that enabled successful navigation, we analyzed the effect of each possible pairing of variables on navigational performance. For each pair of sensory and/or environmental variables, including pairings of a variable with itself, we constructed a list of all tested combinations of values for the two variables. For each combination, we then computed the average path completion across all trials where the variables were set to those values. The results are shown in Figure 5. We chose path completion as the metric of success for this analysis because it represents how close the agent came to successfully navigating the path. A trial in which the agent deviated somewhat from the path for short periods, moving alongside and then quickly recapturing it, is no less successful than one where the agent stayed perfectly on the path. Such small deviations could even represent a more robust set of parameters that enable a broader area in which the agent can correctly identify familiarity. Similarly, an agent that started at an offset but quickly captured the path is just as successful, if not more so, than a similar one that started on the path. Path completion reflects this reasoning.

We expected a number of the trends observed in Figure 5. (The significance of the trends is confirmed by the confidence intervals in Figure 7 in S1 Appendix.) For example, performance improved as the saccade width narrowed and presented the agent with fewer opportunities to get lost. In addition, when navigating based on chemistry alone (*β* = 1.0), a smaller chemical scale (chemistry scale = 0.5mm) is strongly preferred, confirming that small-scale detail of some kind is necessary for successful NFLS.

Less expected, however, was the “U” shaped curve of completion as a function of the number of texture levels. Performance was worse for a sensor with four allowed intensity levels than for a binary one but increased again as the number of levels grew beyond four. We posit that the unexpectedly good performance for a binary sensor could be due to the “smoothing” effect it had on the landscape: Neighboring pixels likely had similar enough values to round to the same binary value. This would cause nearby glimpses to be more similar to one another than they otherwise would be. As a result a binary sensor would be much more forgiving to small deviations from the training path, increasing success rates. The smoothing effect could also explain why the best performance for a binary sensor was seen at the low resolution of 40 × 2, a setting that also blurs the landscape by averaging more landscape pixels into a single sensor pixel.

Binary sensors were also the only configuration where performance was enhanced by increasing the weighting of chemical information in familiarity calculations: performance peaked at an equal *β* = 0.5 balance between textural and chemical information. Surprisingly, sensors with more texture levels performed best when chemical information was neglected (*β* = 0), though performance did not dramatically drop until high weightings of *β* = 0.75 in favor of chemistry were reached. Again surprisingly, the number of chemicals in the environment had no effect on navigational success in our simulations.

### More detail is not always better

The above results suggest, perhaps counter-intuitively, that more detail is not always better for NFLS: The local peak in performance at binary texture resolution, lack of effect from changing the number of chemicals, and small effect of introducing chemical information into the familiarity scores all contradict a simplistic relationship between the number of possible distinct sensor matrices and success. While increasing sensor resolution does increase performance, the effect diminishes to nothing at the transition from a 40 × 4 to a 40 × 6 sensor. The broad range of numbers of texture levels among the most successful sensory configurations (Table 1), despite the clear overall trend of performance increasing as the number of levels increases beyond four (Figure 5), also argues against a direct correlation between performance and the amount of information that can be encoded by the sensor. Similarly, the limited effect on performance of nonzero chemistry weights seemingly contradicts the prevalence of high chemical weights among the best sensory-environmental configurations (Table 3). The most successful *sensory* configurations, however, do *not* have high chemistry weights, indicating that high *β*s are useful only in conjunction with appropriate environmental configurations.

Some of these seeming contradictions could result from scale. It is possible that a high number of texture levels is universally optimal but is not necessary for navigation at the scales considered in this work. It is also possible that in much larger landscapes, where it is necessary to distinguish between many more glimpses, having more chemical information could become unambiguously critical for navigational success.

We propose, however, that these results can be explained through a fundamental interplay in familiarity navigation between “complexity” — the need to distinguish between glimpses in different places and orientations — and “continuity” — the need for small offsets to minimally affect familiarity [37, 52]. Without complexity, the agent cannot choose a direction, and without continuity, the entire navigational scheme is unstable. There exists a degree of intrinsic conflict between complexity and continuity. A white noise landscape is maximally complex but has very low continuity: two identically shaped patches cropped from the landscape, the second offset by a single pixel from the first, are completely uncorrelated. A uniform blank landscape is maximally continuous but has no complexity at all since every glimpse is identical. Less extreme examples can also be constructed. Consider a landscape generated by randomly arranging some number of Gaussians with equal widths. If the width of the Gaussians is small compared to the size of the sensor, continuity will be low but the landscape will have meaningful complexity. As their widths increase, offsets on the scale of the sensor become less and less meaningful, which increases continuity, while the landscape — again at relevant scales — begins to look locally constant, which reduces complexity.

This trade-off could explain many of the observed effects such as the lack of an improvement in performance moving from a 40 × 4 to a 40 × 6 sensor: The larger sensor clearly increases complexity, but it also reduces continuity by better resolving spatial details through smaller sensor pixels. A chemistry weight of *β* = 0.75 dramatically increases continuity, neglecting small-scale textural detail, but in doing so it reduces complexity. The large drop in average path completion from *β* = 0.5 to *β* = 0.75 can be explained as a consequence of the resulting imbalance between complexity and continuity. In the case of *β* = 1.0, texture is completely ignored and the imbalance is even more pronounced, causing navigation to become essentially impossible (see Figure 5).

We believe that this trade-off is a critical area for further research in familiarity navigation, both through large-scale computational simulations and more abstract information theory work. Such work would also be important for the study of non-local (i.e. visual) familiarity. Though visual sensors are inherently highly continuous due to the nature of light, the importance of the relationship between complexity and continuity can still easily be seen. A landscape with many small plants or rocks has high complexity but relatively low continuity because a small change in position or orientation can cause some objects to obscure others, meaningfully changing the scene. A barren landscape with a single large tree in the distance, on the other hand, has very high continuity but lacks complexity.

### Future work

Future work could include the use of biologically inspired neural networks for computing familiarity scores; the explicit simulation of training walks; and the implementation of a step size that dynamically adapts to the overall familiarity of the agent’s immediate environment. Future simulations should also test whether systematic search patterns allow the agent to find the path when it gets lost, as discussed above. Geological and ecological data paired with simplified physical modeling of chemical dispersion and sand deposition could also directly generate more realistic chemo-textural landscapes.

Another valuable question for future work is what mechanism scorpions could use to acquire training glimpses. We note that NFLS, unlike visual familiarity navigation, could function without dedicated training walks. If an agent acquired glimpses while walking away from its burrow, it could in principle construct a set of training glimpses for recapitulating that same path in the opposite direction by flipping those sensor matrices both horizontally and vertically. While there is no biological evidence at present to support such a hypothesis, we believe it deserves attention and could be relevant for robotic applications regardless of its biological plausibility.

We also propose that our analysis of the effects of different parameters on navigational performance, although constrained in this work by scorpion physiology, could serve as a guide for the design of robots that use NFLS, which could have significant advantages for simple navigation tasks such as exiting a room into which a robot had been led. NFLS does not require the unblocked outward view of visual familiarity, the complex algorithms and processing power of machine learning approaches, or the fragility of dead reckoning. Using our model and methodology, roboticists could test parameters relevant to their design choices and extract both optimal configurations and general trends.

## Conclusion

We come to three main conclusions: first, that NFLS is informationally possible using a biologically plausible model of the scorpion’s pectines within a physically plausible model of their environment. More generally, we conclude that familiarity navigation is possible with a local non-visual sensor and that the possibilities for chemo-textural familiarity navigation in scorpions and other animals with complex non-visual sense organs merit further study.

Second, our data strongly suggest a complex relationship, contrary to intuitive expectations, between the informational potential of an NFLS sensor and navigational success. We propose that this relationship rests on the interplay between complexity and continuity and that this result and proposal merit further study as well, both through simulations and information theoretic approaches.

Third, we show that high-throughput simulations — which are novel in the navigation literature to the best of our knowledge — are a powerful tool for assessing behavioral models and their dependence on their parameters. The presence of fragile configurations in our data underlines the importance of testing configurations across many situations using high-throughput methods, even if those situations are quite similar. We believe high-throughput simulation can become an important tool for forming robust conclusions about any behavioral hypothesis that is amenable to simulation.

## Supporting information

**S1 Appendix. Technical details and plots**. Details on: generation of landscapes; generation of synthetic chemical information; estimation of step properties from experimental data; generation of figures; and the implementation of the model.

Discussion of the validity of step limits. Additional result figures.

**S1 Video. Representative trials**. Videos showing a number of representative successes and failures from our simulations.

**S1 File. High-throughput simulation data**. Full results of the high-throughput simulations; parameter files for running them.

**S2 File. Landscape images**. The four landscapes used in this work; the sand images used to generate them.

**S3 File. Motion tracking data**. Scorpion motion tracking data based on [60] used in S1 Appendix to estimate step properties.

## Acknowledgments

A.M. thanks Nili Abrahamsson for her sharp insights, useful discussions, stylistic feedback, and ongoing support.

We also thank Mariëlle Hoefnagels for carefully reviewing the manuscript and her many valuable suggestions for improvement.

The authors thank the other members of the Gaffin Scorpion Lab — Tanner Ortery, Safra Shakir, Drew Doak, and Jacob Sims — for their feedback and discussions on early work.

The computing for this project was performed at the OU Supercomputing Center for Education and Research (OSCER) at the University of Oklahoma (OU). The authors gratefully acknowledge the staff of the center for their helpfulness.

